# P-DOR, an easy-to-use pipeline to reconstruct outbreaks using pathogen genomics

**DOI:** 10.1101/2023.05.30.542810

**Authors:** Gherard Batisti Biffignandi, Greta Bellinzona, Greta Petazzoni, Davide Sassera, Gian Vincenzo Zuccotti, Claudio Bandi, Fausto Baldanti, Francesco Comandatore, Stefano Gaiarsa

## Abstract

Bacterial Healthcare Associated Infections (HAIs) are a major threat worldwide, which can be counteracted by establishing effective infection control measures, guided by constant surveillance and timely epidemiological investigations. Genomics is crucial in modern epidemiology but lacks standard methods and user-friendly software, accessible to users without a strong bioinformatics proficiency. To overcome these issues we developed P-DOR, a novel tool for rapid bacterial outbreak characterization. P-DOR accepts genome assemblies as input, it automatically selects a background of publicly available genomes using k-mer distances and adds it to the analysis dataset before inferring a SNP-based phylogeny. Epidemiological clusters are identified considering the phylogenetic tree topology and SNP distances. By analyzing the SNP-distance distribution, the user can gauge the correct threshold. Patient metadata can be inputted as well, to provide a spatio-temporal representation of the outbreak. The entire pipeline is fast and scalable and can be also run on low-end computers.

**Availability and implementation:** P-DOR is implemented in Python3 and R and can be installed using conda environments. It is available from GitHub https://github.com/SteMIDIfactory/P-DOR under the GPL-3.0 license.

## INTRODUCTION

Bacterial infections are a constant threat to public health worldwide. When dealing with Healthcare-Associated Infections (HAIs) and outbreaks, timely epidemiological investigation is pivotal to establish effective infection control measures (Jiang *et al*., 2015; Balloux *et al*., 2018; Harris *et al*., 2013; Raven *et al*., 2017). Despite their wide use, conventional molecular typing techniques, such as Pulsed Field Gel Electrophoresis (PFGE) and Multi-Locus Sequence Typing (MLST), have a lower discriminatory capability in comparison to the modern Whole Genome Sequencing (WGS)-based typing, while maintaining similar costs and timescales.

Over the past few years, WGS-based typing has been increasingly adopted, first for research purposes and then as a routinary screening tool for infectious disease epidemiology in hospitals and public health settings. This approach leverages *in silico* techniques for isolates typing, antimicrobial profile determination, and outbreak reconstruction (Jiang *et al*., 2015; Balloux *et al*., 2018; Harris *et al*., 2013; Raven *et al*., 2017; Sherry *et al*., 2019; Onori *et al*., 2015; Ferrari *et al*., 2019). Several computational methods to analyze such datasets have been developed, which include database design (Lam *et al*., 2021; Zhou *et al*., 2020), epidemiological models via Bayesian inference (Campbell *et al*., 2018; De Maio *et al*., 2016; Jombart *et al*., 2014), network analysis (Worby *et al*., 2017, 2014) and phylogeny (Didelot *et al*., 2021).

Relationships among strains are mainly inferred using Single-Nucleotide Polymorphisms (SNPs) or k-mers. When reconstructing outbreaks, strains isolated from different sources (e.g. patients, fomites) and having SNP-distances below specific thresholds can be considered part of the same transmission cluster. The network of these genetically correlated strains can be used to reconstruct the pathogen transmission route. Although threshold-based methods are largely applied in genomic epidemiology (Hatherell *et al*., 2016; Octavia *et al*., 2015; Dallman *et al*., 2015; David *et al*., 2019), they lack standardization (Duval *et al*., 2023). Indeed, threshold values can vary across bacterial species/clones because of their different genomic architectures (e.g. mutation rate, recombination). Also the duration of the epidemic event analyzed can influence the genetic variability in the bacterial population: a SNP-distance threshold set to disentangle a short outbreak can be inappropriate for a long-term genomic surveillance study (Duval *et al*., 2023). Furthermore, SNP distances can be affected (even by tenths or hundreds) by the SNP calling approach (e.g. mapping reads or aligning assembled contigs) and by the reference genome and software used. Finally, the sole use of genomic data without the inclusion of other information like clinical metadata (e.g. sample date/type, hospitalization ward) limits the comprehension of epidemic events (Stimson *et al*., 2019; De Maio *et al*., 2016; Jombart *et al*., 2014; Didelot *et al*., 2021; Duval *et al*., 2023).

Most of the software available for WGS-based epidemiological investigation is not user-friendly (De Maio *et al*., 2016; Zhou *et al*., 2020; Didelot *et al*., 2021; Campbell *et al*., 2018), not free (e.g. SeqSphere+ Ridom GmbH software), and/or does not encompass all the analyses required for a comprehensive study (De Maio *et al*., 2016; Zhou *et al*., 2020; Didelot *et al*., 2021). Most of the methods require the user to have a computational background, as they are composed of multiple command-line tasks that must be serially performed in succession, and often require format changes. This prevents most clinicians from performing genomic investigations in first person and limits their understanding of the results. Consequently, it also hampers them from making epidemiological conclusions in light of both clinical information and of their past experience on the field, which in turn would enable them to provide valuable feedback to developers. On the other hand, online tools are available, which are accessible to a wider usership, but lack the tunability that is required for most epidemiological investigations (e.g. (Trifinopoulos *et al*., 2016)) and are restricted to single tasks (e.g. phylogeny).

To answer the need for a comprehensive, tunable, and user-friendly tool, we developed P-DOR, a bioinformatic pipeline for rapid WGS-based bacterial outbreak detection and characterization. P-DOR integrates genomics and clinical metadata and uses a curated global genomic database to contextualize the strains of interest within the appropriate evolutionary frame. P-DOR is available at https://github.com/SteMIDIfactory/P-DOR.

## P-DOR WORKFLOW

The inputs for the core P-DOR analysis are: i) a folder containing the query genome assemblies; ii) a reference genome for SNP extraction; iii) a sketch database file generated by Mash (Ondov *et al*., 2016); iv) a table containing the patient metadata (i.e. hospitalization ward, date of admission and discharge). This last input is not mandatory, but when provided, it will be integrated in the analysis to add further clues on the epidemic event. The query genomes of the study must be in FASTA format, and can be complete or draft assemblies.

Sketch files contain the genomic information of the strains from a Source Dataset (SD) chosen by the user. A sketch file is a vastly reduced representation of the genomes, which is produced via the *MinHash* algorithm to allow fast distance estimation using low memory and storage requirements. Regularly updated sketches for each of the ESKAPE members (*Enterococcus faecium, Staphylococcus aureus, Klebsiella pneumoniae, Acinetobacter baumannii, Pseudomonas aeruginosa* and *Enterobacter* spp.) are available in the P-DOR repository. Personalized SD sketches can alternatively be built by the user using the “makepdordb.py” script. This script can automatically download the high-quality genomes of a species from the BV-BRC collection (Davis *et al*., 2020) or build a custom SD sketch starting from any set of genomes. The sketch files are used to compute the k-mer distances between each query genome and the SD genomes. Then, for each query genome the *n* most similar SD genomes are selected and joined in a Background Dataset (BD). Lastly, the query genomes are joined with the BD to obtain the Analysis Dataset (AD). Optionally, the entire AD can be scanned for the presence of antimicrobial resistance and virulence genes using AMRFinderPlus (Feldgarden *et al*., 2021).

After that, each genome of the resulting AD is aligned to the reference genome using Nucmer, the alignment obtained is polished using Delta-Filter and SNPs are called using show-snps, all the commands being part of the Mummer4 package (Marçais *et al*., 2018). Lastly, the SNPs of all AD genomes are combined into a coreSNPs alignment using a Python script (Ferrari *et al*., 2019). Then, the coreSNPs alignment is used to infer a Maximum Likelihood (ML) phylogeny via the IQ-TREE (Nguyen *et al*., 2015) software, which includes a prior step for the selection of the substitution model.

To infer relationships among the strains, epidemiological clusters are inferred on the basis of coreSNPs distances using a threshold value. The user can manually set the SNP threshold parameter according to previous studies in the literature, or by visualizing the SNP pairwise distances distribution plot provided among the outputs of P-DOR. The user should analyze the results, possibly tweak the threshold parameter and run P-DOR again.

## OUTPUTS

The main outputs of P-DOR are: i) a SNP-based phylogenetic tree; ii) a heatmap reporting the phylogenetic tree and presence/absence of resistance and virulence factors; iii) a heatmap showing the coreSNP distance matrix; iv) the histogram of the distribution of the SNP distances; v) a graph visualization of the epidemiological clusters, in which each pair of strains (nodes) is connected if the coreSNP distance between them is below the threshold. The epidemiological clusters are also highlighted on the phylogenetic tree. In addition, if patients metadata are provided, P-DOR creates a spatio-temporal representation of the outbreak and outputs a patients timeline plot where strains are placed based on the date of isolation and are connected on the basis of epidemic clusters.

## PERFORMANCE TEST

To test P-DOR, we simulated the genomic sequences of 11 *Klebsiella pneumoniae* isolates, involved in a complex epidemic event, with two distinct bacterial strains (both belonging to Sequence Type 258) circulating in a hospital in the same period. In detail, we obtained six simulated sequences starting from genome NJST258_1 and 5 sequences from genome NJST258_2 using the software simuG (Yue and Liti, 2019). The genomes were simulated in a hierarchical way: i.e. each generated genome differs one to five SNPs from the parent. The simulated dataset was used as query to P-DOR. The SD was obtained from BV-BRC on 16 April 2023 using the script makepdordb.py. The Background Dataset (BD) was built selecting the 20 closest SD genomes for each query genome, as described above. After analyzing the distribution of SNP distances, the threshold was set at 21 SNPs (Figure 1A). Finally, the complete genome of strain HS11286 (NZ_CP029384.2) was used as a reference for SNP calling. The phylogeny obtained correctly determined the presence of two outbreaking strains (Figure 1B). P-DOR divided the simulated genomes into two monophyletic clusters labeled C1 (green) and C4 (orange). Both clusters also include a background genome (BV-BRC codes 1420013.3 and 1420012.3), which correspond to isolates NJST258_2 and NJST258_1 i.e. the genomes used as starting points for the generation of the simulated sequences. These results demonstrate the capability of the P-DOR pipeline to select a background apt for epidemiological investigations and to identify outbreak clusters. They also show that P-DOR can identify the putative source of each epidemic cluster, when it is available in the SD. Figure S1 shows the SNP distances among all genomes in analysis and further confirms the results observed in the phylogeny (Figure 1B). Furthermore, the timeline (Figure 1C) can be used to hypothesize the chain of transmission based on the dates of isolation and the epidemiologic classification of the isolates.

**Figure 1.**
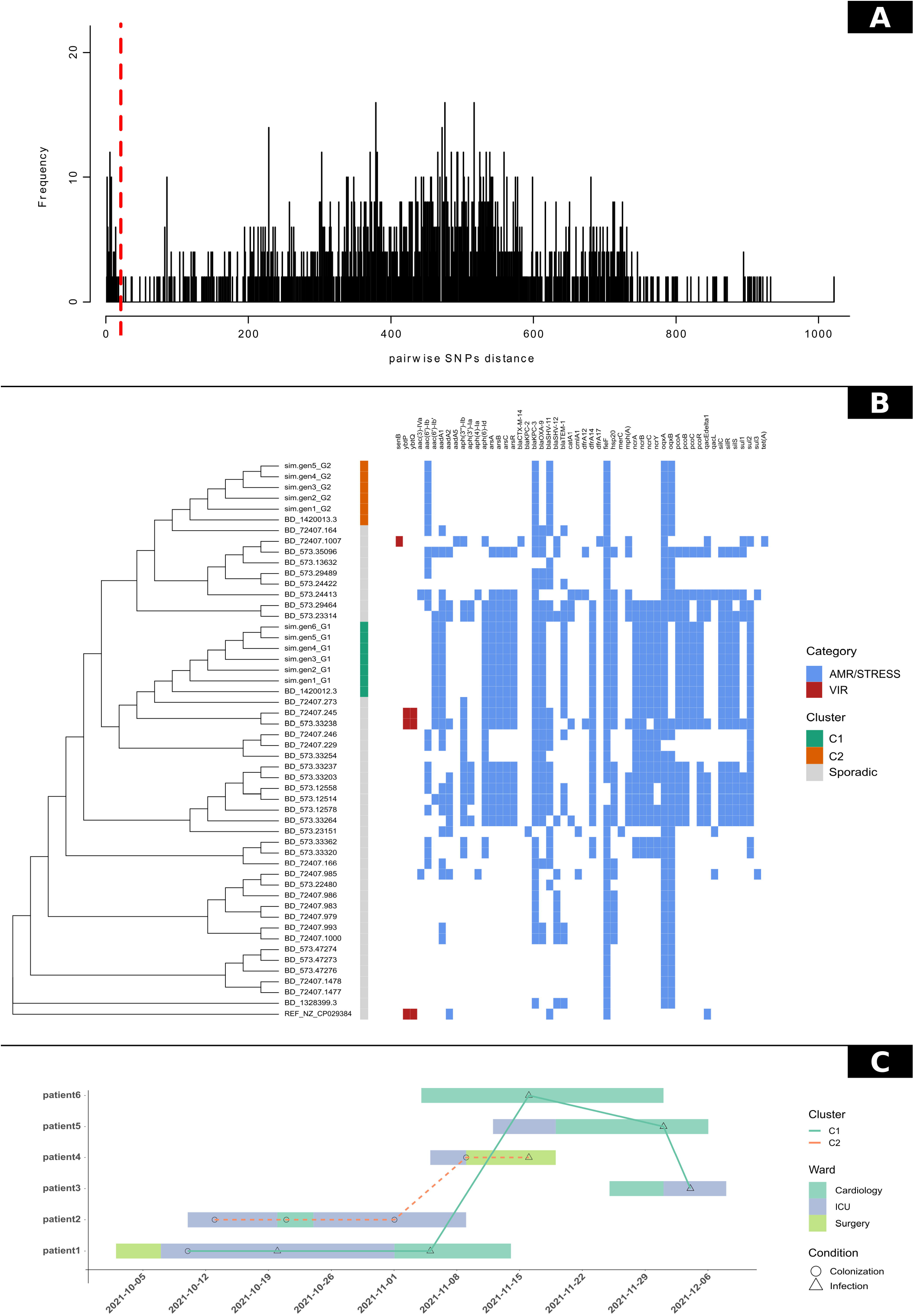
The P-DOR outputs of the test analysis. A) Phylogenetic tree of the Analysis Dataset (AD). The first column shows epidemiological clusters of strains with SNP distances below the threshold set by the user. In addition, a heatmap representing the detection of resistance (blue) and virulence (orange) determinants is shown next to the tree for a better representation of the epidemic event. B) Distribution of SNP distances calculated between all permutations of genome pairs in the AD. C) Timeline of the patients movements during the hospitalization. Points indicate outbreak genomes and are shaped according to the isolation source of the corresponding strain. Samples are linked if their genetic distance in terms of SNPs is below the threshold.

The analysis was performed on a total of 50 genomes using a maximum of 313 Mb of RAM. The entire process required one hour and 37 minutes, using four threads at 2.3 Ghz, 29 minutes when using 20 threads; these numbers drop respectively to 2.9 and 2.1 minutes when excluding the time-consuming AMRFinderPlus step (default setting). These results show that P-DOR is a fast tool, which can be used in clinical contexts even when high informatic skills or resources are not available. Future efforts will be focused on developing a web-based interface, to further improve the ease of use and removing the need for computational resources.

## FIGURE LEGENDS

**Figure S1.** Heatmap representing the pairwise SNPs distance between all pairs of genomes in the test analysis.

