## Supplementary figures and images for "P-DOR, an easy-to-use pipeline to reconstruct outbreaks using pathogen genomics"

### Figure S1

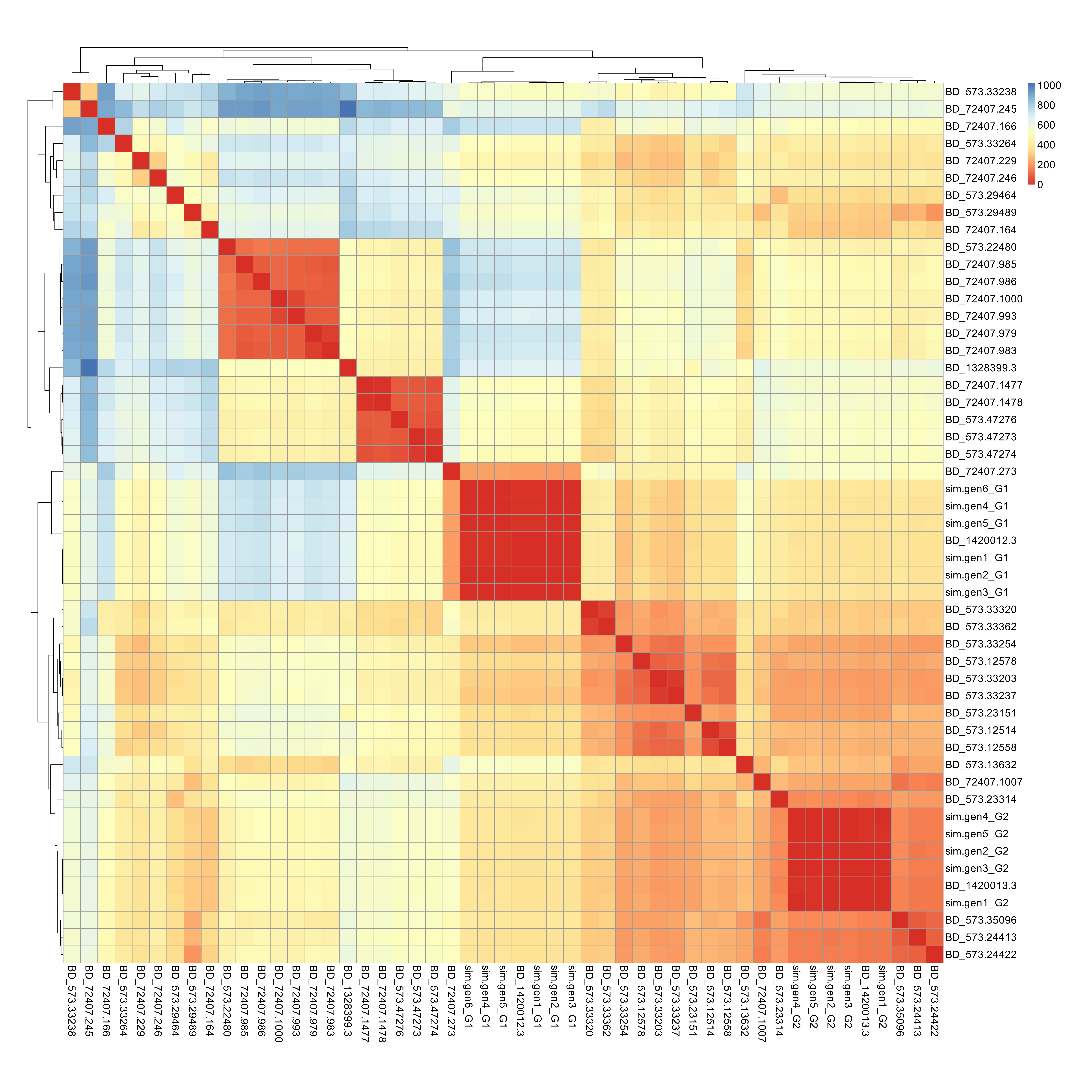
